# J chain dictates the recognition of IgA by FcRL4

**DOI:** 10.1101/2023.12.18.572144

**Authors:** Chen Su, Yuxin Wang, Xiaoke Yang, Junyu Xiao

## Abstract

Fc-receptor-like 4 (FcRL4) selectively recognizes systemic IgA. The cryo-electron microscopy structure of the FcRL4–Fcα (IgA-Fc)–Joining (J) chain complex reveals that FcRL4 interacts with Fcα–J in a 1:1 stoichiometry, with FcRL4 primarily interacting with the J chain. The binding of Fcα–J to FcRL4 would hinder the binding of pIgR, but not FcαRI. These findings provide new insights into IgA and emphasize significant differences between various IgA receptors.

---

Immunoglobulin A (IgA) plays a crucial role in the human immune system as the primary antibody on mucosal surfaces and also the second most abundant antibody in serum. IgA can exist in monomeric or polymeric forms, with the latter consisting of two to five monomeric IgAs linked together by the joining chain (J chain)^1,2^. The J chain-containing IgA polymers can be further transported to mucosal surfaces by the polymeric immunoglobulin receptor (pIgR). The extracellular region of pIgR, known as the secretory component or SC, associates with the transported IgA to form secretory IgA (SIgA). In addition to pIgR, IgA can interact with several other Fc receptors, including the FcαRI (CD89) and FcαμR (CD351)^3^. Moreover, recent evidence has demonstrated that FcRL4 is a newly identified member of the IgA receptor family that specifically recognizes systemic IgA^4,5^.

FcRL4 belongs to the Fc-receptor-like (FcRL) family, which includes six cell surface proteins (FCRL1-6, as shown in Fig. 1a), as well as two intracellular proteins located in the endoplasmic reticulum (FCRLA and FCRLB)^6^. These FcRL proteins are characterized by multiple Ig extracellular domains and immunoreceptor tyrosine-based activation motif (ITAM) and/or immunoreceptor tyrosine-based inhibition motif (ITIM) in their intracellular domain, enabling them to modulate cellular and humoral immunity^7,8^. In contrast to the Fc receptors primarily expressed by myeloid cells^9^, FcRLs are predominantly expressed by lymphocytes, particularly B cells. For example, FcRL4 is expressed on a subset of memory B cells found in mucosal-associated lymphoid tissues such as tonsils and Peyer’s patches^10,11^. FcRL4 contains three ITIM sequences in its intracellular domain^12^, suggesting its likely function as an inhibitory receptor for IgA^13,14^. Additionally, FcRL4 has been implicated in autoimmune diseases. In rheumatoid arthritis, the FcRL4^+^ memory B cells are identified as a pro-inflammatory cell population in synovial fluid, potentially contributing to chronic inflammation and bone destruction^15,16^. However, the molecular mechanism underlying FcRL4 function remains incompletely understood.

**Figure 1.**
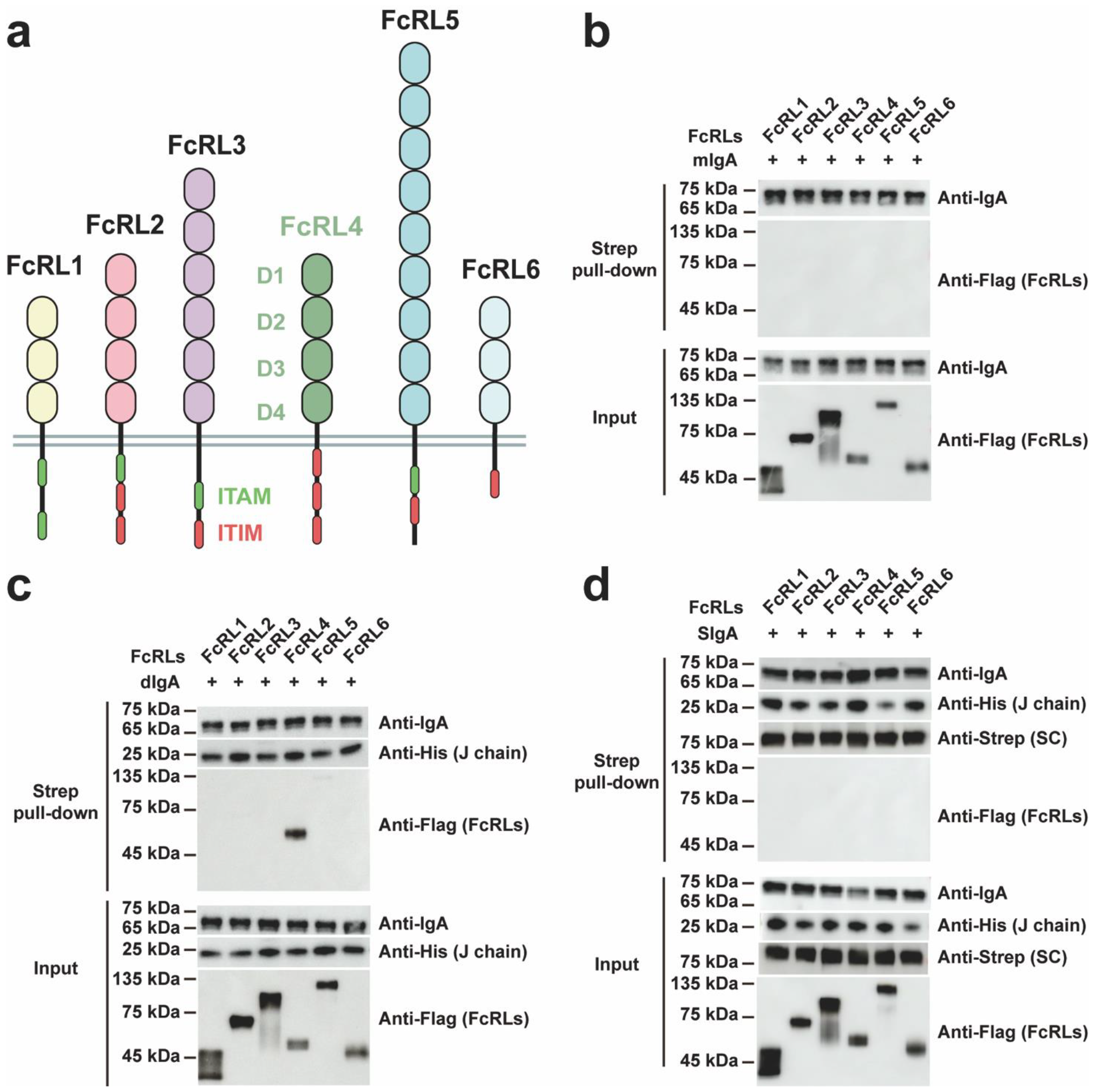
FcRL4 specifically recognizes systemic IgA. **a**. Domain organizations of FcRL1–6 proteins. **b**. None of the FcRL proteins bind to monomeric IgA. The purified mIgA-604 was pre-incubated with Flag-tagged FcRLs and was subject to immunoprecipitation using Capto-L beads. The result was analyzed by SDS-PAGE and western blot analysis using anti-Flag tag and anti-IgA antibodies. **c**. FcRL4, but not by other FcRLs, specifically interacts with dIgA-604. The His-tagged J chain was detected with anti-His antibody. **d**. None of the FcRL proteins bind to SIgA-604. The Strep-tagged SC was detected with anti-Strep antibody.

Using cell lines expressing FcRL4, Wilson et al. initially observed the binding of heat-aggregated IgA to FcRL4^4^. Subsequently, Liu et al. demonstrated that FcRL4 recognizes systemic IgA linked by J chain in the absence of heat aggregation, but not monomeric IgA^5^. To confirm the direct interaction between FcRL4 and IgA, we expressed and purified Flag-tagged extracellular domains of human FcRL1–6 (Extended Data Fig. 1a). We also generated a monomeric IgA antibody, mIgA-604, which contains a heavy chain engineered by conjugating the VH domain of BD-604, a SARS-CoV-2 neutralizing IgG antibody^17^, to the constant region of human IgA1, and the light chain of BD-604. This IgA is co-expressed with the J chain and secretory component to produce the dimeric and secretory forms (dIgA-604 and SIgA-604), respectively (Extended Data Fig. 1a). Protein binding studies using these purified components revealed that none of the FcRL proteins interacted with mIgA-604 (Fig. 1b), while FcRL4 specifically bound to the J chain-containing dIgA-604 (Fig. 1c). Furthermore, the presence of secretory component fully blocked the interaction between SIgA-604 and FcRL4 (Fig. 1d), in agreement with previous analyses^5^. These biochemical results validate previous findings obtained using cell systems and unequivocally demonstrate the specific interaction between FcRL4 and dIgA. It is noteworthy that, although a recent study has reported the binding of FcRL3 to SIgA^18^, we did not observe direct binding between the extracellular region of FcRL3 and SIgA-604 (Fig. 1d).

To further investigate the FcRL4–dIgA interaction and compare it with the binding affinities of IgA with other IgA receptors, we engineered the extracellular regions of FcRL4, FcαRI, and pIgR (aka SC) with C-terminal Twin-Strep tags. These proteins were then immobilized on biosensor chips coated with covalently attached StrepTactin XT, an engineered streptavidin with picomolar affinity to the Twin-Strep tag, to mimic the orientation of these receptors on the cell surface. As a control, we also engineered green fluorescent protein (GFP) in a similar manner. Subsequently, we conducted surface plasmon resonance (SPR) analyses using Fcα–J, the dIgA core consisting of an Fcα (IgA-Fc) dimer and the J chain, as the analyte. Our results revealed that FcRL4 exhibited a 5 nM affinity for Fcα–J (Fig. 2a, Extended Data Fig. 1b), which is comparable to the high-affinity interaction observed between FcαRI and Fcα–J (Fig. 2b). It is important to note that our measured affinity of FcαRI–Fcα–J is consistent with previous findings^19^. Additionally, as FcRL4 possesses only a single binding site on Fcα–J (see below), its affinity for Fcα–J is arguably higher than that of FcαRI, as the bivalent interaction of Fcα–J with two surface-anchored FcαRI^20^ would lead to higher avidity and, therefore, a higher apparent binding affinity. Furthermore, the affinity of FcRL4 for Fcα– J was found to be higher than that of pIgR (Kd of 23 nM, Fig. 2c), which also has a single binding site on Fcα–J.

**Figure 2.**
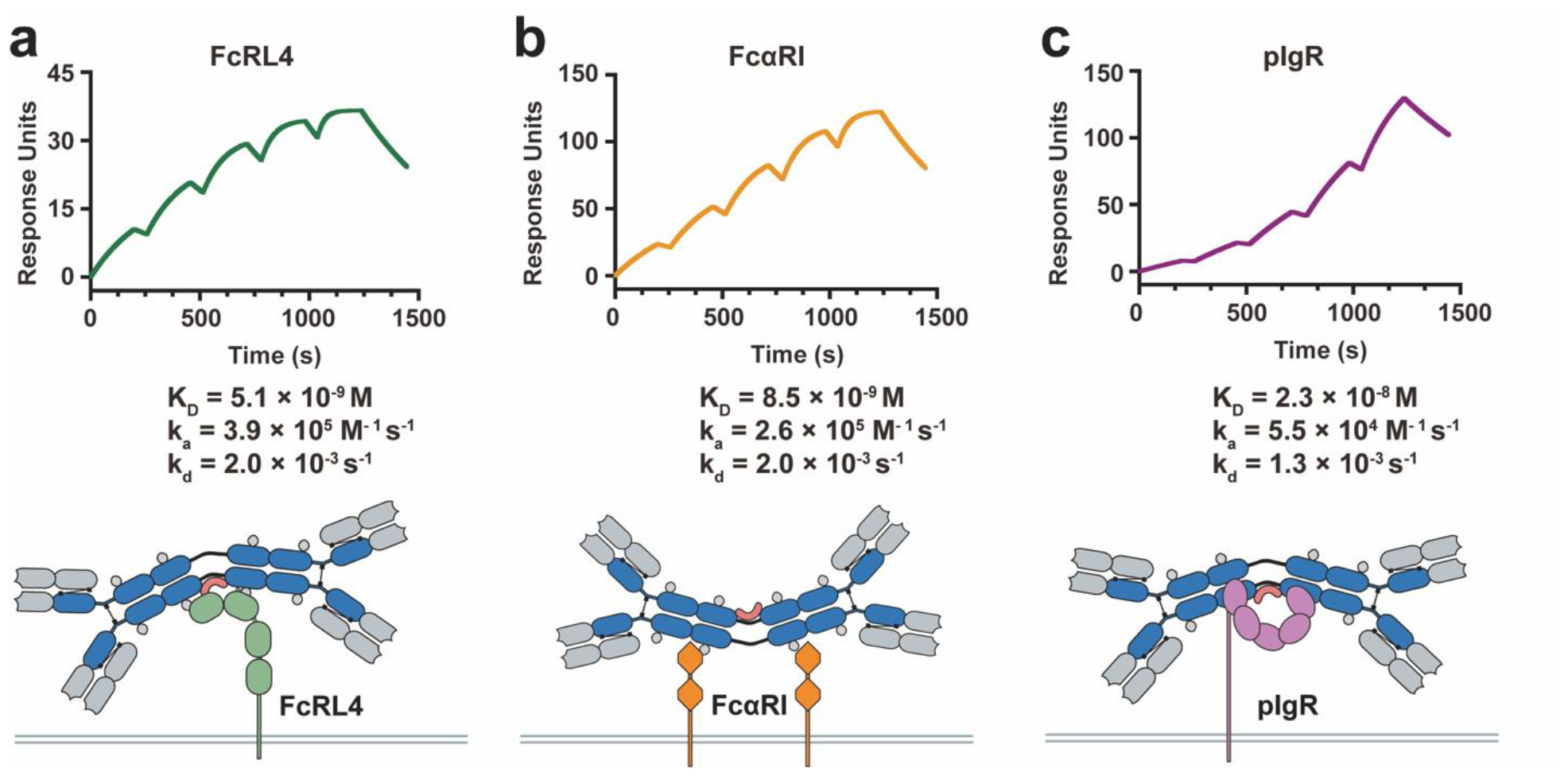
FcRL4 binds to systemic IgA with a high affinity. **a**. Single cycle kinetics SPR analysis of the interaction between Fcα–J and FcRL4. The extracellular region of FcRL4 with a C-terminal twin-strep tag was captured on the chip coated with covalently attached StrepTactin XT. Fcα–J at 5, 10, 20, 40, and 80 nM was then passed sequentially through the chip. All SPR experiments in this paper have been repeated at least three times. **b**. Single cycle kinetics SPR analysis of the interaction between Fcα–J and FcαRI. The extracellular region of FcαRI with a C-terminal twin-strep tag was immobilized as above. **c**. Single cycle kinetics SPR analysis of the interaction between Fcα–J and immobilized pIgR.

To elucidate the molecular mechanism of FcRL4–dIgA interaction, we conducted cryo-electron microscopy (cryo-EM) analysis of the FcRL4–Fcα–J complex (Extended Data Fig. 1c). Data were collected using a tilt strategy to resolve issues with preferred sample orientation, and the FcRL4–Fcα–J structure was determined at a resolution of 3.3 Å (Fig. 3a, Extended Data Fig. 2, Extended Data Table 1). The overall structure of Fcα–J is similar with Fcα–J seen in SIgA, with two Fcα molecules held together by the J chain in a bent fashion. A single FcRL4 binds to Fcα–J, revealing a 1:1 FcRL4:Fcα–J binding stoichiometry that is distinct from the 2:1 FcαRI:Fcα–J interaction^21^.

**Figure 3.**
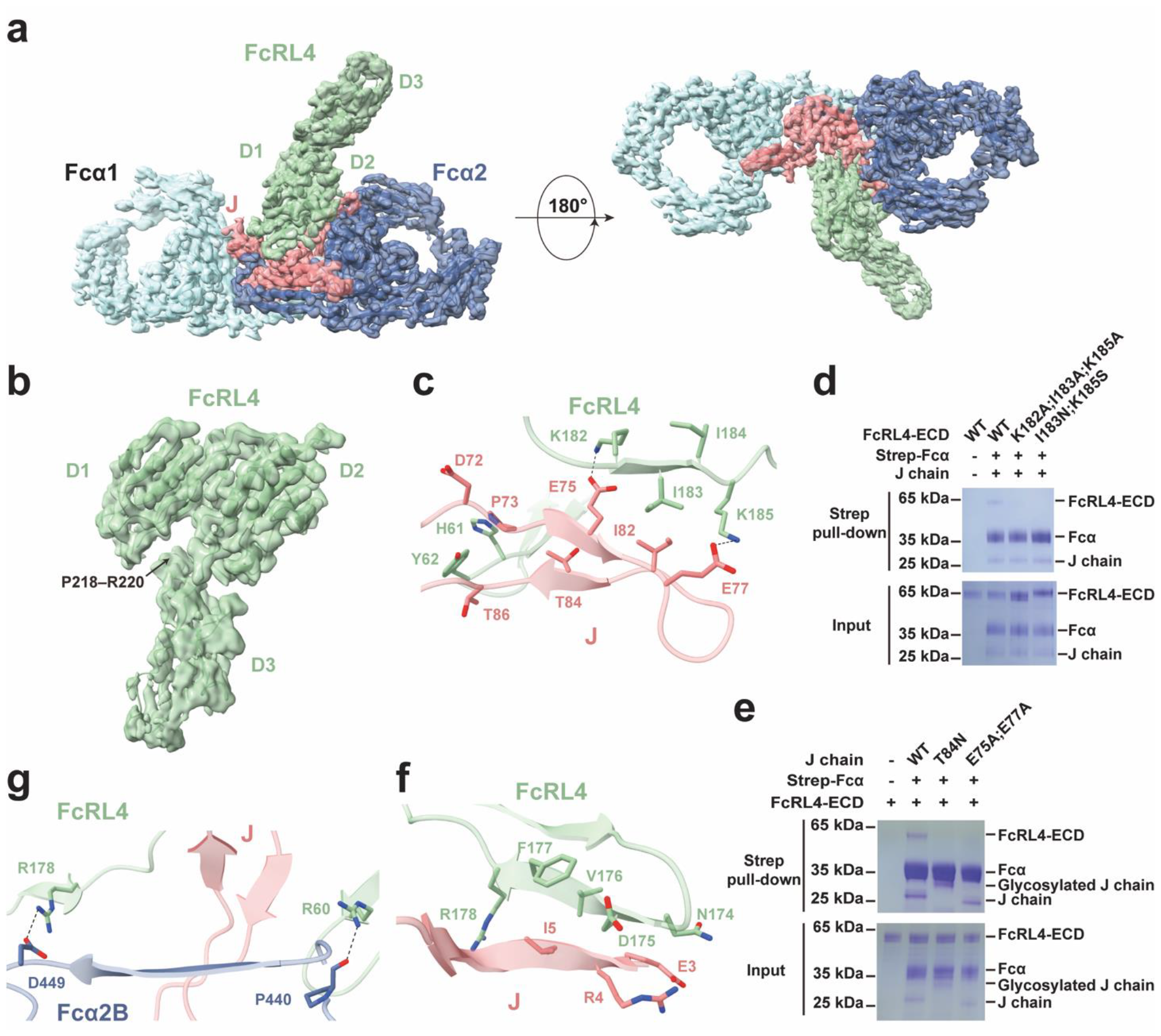
The cryo-EM structure of the Fcα–J–FcRL4 complex. **a**. The cryo-EM structure of the human Fcα–J–FcRL4 complex is shown in two views. Fcα1 and Fcα2 are shown in light cyan and blue, respectively. J chain is shown in light coral, and FcRL4 is shown in green. Similar color schemes are used in all figures unless otherwise indicated. **b**. Structure of FcRL4. **c**. Interactions between FcRL4 and the β5–β6 hairpin of J chain. **d**. The FcRL4 K182A;I183A;K185A mutation displayed reduced interaction with Fcα–J, and the FcRL4 I183N;K185S abolished interaction with Fcα–J in a pull-down assay. **e**. Fcα–J containing J chain mutants T84N or E75A/E77A exhibited a diminished interaction with FcRL4. **f**. Interactions between FcRL4 and the N-terminal region of the J chain. **g**. Interactions between FcRL4 and Fcα2.

The extracellular region of FcRL4 comprises four immunoglobulin-like domains (D1–D4). The D1–D2 domains and the N-terminal portion of D3 exhibit high-quality EM densities and can be distinctly resolved. The overall assembly of D1–D3 presents a ‘7’-like appearance (Fig. 3b). Specifically, D1 and D2 tightly pack together in an antiparallel manner. Furthermore, D3 is positioned beneath D1–D2, with residues Pro218– Arg220 wedging into the interface between D1 and D2.

The J chain plays a critical role in determining the binding of FcRL4. A groove situated between the first and second domains of FcRL4 provides a space for the β5–β6 hairpin of the J chain. Within this groove, Tyr62 is positioned between Asp72_J_–Pro73_J_ and Thr86_J_, and Ile183 interacts with Ile82_J_ (Fig. 3c). Additionally, Lys182 forms a salt bridge with Glu75_J_, while Lys185 contacts Glu77_J_. Two FcRL4 mutants, K182A/I183A/K185A and I183N/K185S, were used to disrupt the interactions between FcRL4 and Fcα–J, confirming the functional significance of these interactions (Fig. 3d). Furthermore, two Fcα–J variants carrying J chain mutants, T84N and E75A/E77A, with unchanged structural integrities, exhibited reduced interactions with FcRL4 (Extended Data Fig. 1d–f, Fig. 3e). Similar to the FcRL4 I183N/K185S mutant, the T84N mutation was designed to create an Asn–X–Ser/Thr consensus sequence for N-linked glycosylation, as Thr84 was followed by Ala85–Thr86 in the J chain.

The D2 domain of FcRL4 also interacts with the N-terminal region of the J chain, involving residues Asn174–Arg178 of FcRL4 and Glu3_J_–Ile5_J_ (Fig. 3f). In fact, this interaction leads to the formation of a short β-strand between Glu3_J_–Ile5_J_. Importantly, FcRL4 also interacts with the Fcα chain, with Arg60 forming a hydrogen bond with the main chain carbonyl group of Pro440_Fcα2B_, and Arg178 making an ionic contact with Asp449_Fcα2B_ (Fig. 3g). Together, these interactions contribute to the highly specific binding between FcRL4 and Fcα–J. Importantly, most of the residues involved in interacting with Fcα–J are unique to FcRL4 and are not present in other FcRL family members (Extended Data Fig. 3), providing a rationale for the unique ability of FcRL4 to recognize J chain-containing IgA (Fig. 1c).

Modeling based on the structures of tetrameric and pentameric IgA indicates that FcRL4 would likely recognize these higher-order IgA polymers as well (Extended Data Fig. 4a, b). However, FcRL4 would not be able to recognize secretory IgA (SIgA) that contains secretory component (SC), as observed in the structural alignment of the FcRL4–Fcα–J complex to the SIgA core, revealing that SC would obstruct the binding of FcRL4 (Extended Data Fig. 4c), consistent with previous analyses^5^. Conversely, modeling based on the structure of IgA in complex with FcαRI suggests that the two FcαRI-binding sites in dIgA remain accessible in the presence of FcRL4 (Extended Data Fig. 4d). This suggests the intriguing possibility that FcRL4 and FcαRI, whether on the same cell or from different cells, could potentially bind systemic IgA simultaneously.

In contrast to the extensively studied IgA Fc receptor FcαRI, FcRL4 is unique for its unique recognition of a composite binding site on systemic IgA, involving both the J chain and Fcα. Our research also confirms that FcRL4 exhibits high-affinity binding to Fcα–J (Kd of 5 nM), consistent with previous evidence indicating the constitutive occupation of FcRL4 on primary memory B cells by systemic IgA^5^. While the full biological function of FcRL4 is yet to be fully elucidated, its unique recognition of J chain-containing systemic IgA, but not monomeric and secretory IgA, endows it with the ability to convey specific signaling events. Notably, mice lack a FcRL4 homolog^6^, similar to their lack of FcαRI, suggesting significant differences in IgA-related immune functions between humans and mice.

## Materials and Methods

### IgA binding assays

For mIgA-604, HEK293 cells were transiently transfected with two plasmids expressing the heavy chain containing V_H_ of BD-604 neutralizing IgG antibody conjugated with IgA1 constant region and the corresponding light chain. dIgA-604 or SIgA-604 were generated by transfection of plasmids encoding heavy and light chains of mIgA-604, as well as the J chain containing an 8-residue His, and with or without SC. mIgA-604 were purified by anti-light chain affinity chromatography using Capto-L beads (GE Healthcare). dIgA-604 and SIgA-604 were purified using Ni-NTA beads (Smart-Lifesciences) and a Superose 6 Increase column (GE Healthcare).

The extracellular domain of FcRL1 (Uniprot Q96LA6, residues 17-307), FcRL2 (Uniprot Q96LA5, residues 20-401), FcRL3 (Uniprot Q96P31, residues 18-573), FcRL4 (Uniprot Q96PJ5, residues 20-387), FcRL5 (Uniprot Q96RD9, residues 16-851) and FcRL6 (Uniprot Q6DN72, residues 20-307) with an 8-residue His tag and a Flag tag at C-terminal were ligated into the modified pcDNA vector with a N-terminal IL-2 signal peptide. These FcRL proteins were purified by Ni-NTA beads as described above.

The IgAs and FcRLs were incubated with the Capto-L beads in the binding buffer (25 mM Tris-HCl, 150 mM NaCl, pH 7.4) and rotated at 4 °C for 1 h. After the incubation, the beads were spun down and washed three times with the binding buffer. The samples were analyzed by SDS-PAGE and detected by immunoblotting using antibodies to the IgA heavy chain (1:3,000; ab124716, Abcam) and antibodies to the Strep tag, His tag and Flag tag (1:1000; F3165, Sigma).

### Surface plasmon resonance

The surface plasmon resonance experiments were performed on the Biacore 8K system (GE Healthcare). The IgA-receptors were surface captured via StrepTactin XT (Twin-Strep-Tag Capture Kit, IBA GmbH) covalently immobilized on a CM5 chip. As a negative control, the GFP was immobilized on the chip similarly. The reference flow cell was left blank. All assays were performed with a running buffer of HBS-EP (10 mM HEPES pH 7.4, 150 mM NaCl, 3 mM EDTA, 0.05% Tween-20) at 25 °C to determine the binding kinetics between the IgA and its receptors. IgA was injected over the flow cell at a range of five concentrations prepared by serial twofold dilutions at a flow rate of 30 μL/min using a single-cycle kinetics program. All data were fitted to a 1:1 binding model using Biacore Insight Evaluation 4.0. Each SPR experiment was repeated at least three times.

### Protein expression and purification for cryo-EM

The extracellular domain (residues 20-387) of human FcRL4 with a C-terminal 8-residue His tag was cloned into a modified pcDNA vector with a N-terminal IL-2 signal peptide. The constructs encoding Fcα and J chain were previously described^22^. The plasmids expressing Fcα–J and FcRL4 was co-transfected into the HEK293F cells by polyethylenimine (Polysciences). The cells were cultured at 37 °C, 5% CO_2_ and 55% humidity in SMM 293T-I medium (Sino Biological Inc.) for 72-96 hours until harvest. The conditioned media were collected by centrifugation, then concentrated using a Hydrosart Ultrafilter (Sartorius), and exchanged into the binding buffer (25 mM Tris-HCl, pH 7.4, 150 mM NaCl). The protein was first isolated by Ni-NTA affinity purification and eluted with the binding buffer supplemented with 500 mM imidazole. The eluted protein was further purified using a Superose 6 increase column (GE Healthcare) pre-equilibrated with the binding buffer, to isolate the Fcα–J–FcRL4 complex.

### Cryo-EM sample preparation and data collection

4 μL of Fcα–J–FcRL4 (0.3 mg/mL) were applied to glow-discharged holy-carbon gold grids (Quantifoil, R1.2/1.3, 300 mesh) or Ni-Ti grids (CryoMatrix, R1.2/1.3, 300 mesh). The grid was blotted at 4 °C, 100% humidity, and flash-frozen into the liquid ethane with a Vitrobot Mark IV (FEI). A Talos Arctica microscope equipped with Ceta camera (FEI) was used to screen grids. Data collection was performed on a 300 kV Titan Krios electron microscope (FEI) operated with a K3 Summit direct electron detector (Gatan) and the Gatan Imaging Filter (GIF) energy-filtering slit width set at 20 eV. Movie stacks were recorded in super-resolution mode at a magnification of 105,000× using the EPU software (E Pluribus Unum, Thermo Scientific), which corresponded to a pixel size of 0.83 Å per pixel. The defocus range was set from –1.1 to –1.5 μm. Each movie stack was recorded for 2.56 s and subdivided into 32 frames with a total electron exposure of 59.5 electrons per Å^2^. The dose rate was 16 e^−^/pixel/s. Statistics for data collection are summarized in Extended Data Table 1.

### Imaging processing, model building and structure refinement

A total of 7263 movies were collected at 0° tilt, 3088 movies were collected at 15° tilt and 2345 movies were collected at 30° tilt from different grids for the structural analysis of Fcα–J–FcRL4 complex. The image processing was performed using cryoSPARC^23^. Raw movies were motion-corrected and dose-weighted with Patch motion correction, and the contrast transfer function (CTF) parameters were estimated by Patch CTF estimation. Particle picking was performed by Blob picker and Template picker to generate a dataset of 3,314,645 particles. Several rounds of 2D classification were performed to remove noise, and 1,215,993 particles were kept for generating a reference 3D map using Ab-initio reconstruction, followed by Heterogeneous Refinement to further clean particles. Among the six 3D classifications, only one showed the complete complex (369,590 particles). The particles in this qualified group were subjected to Homogeneous Refinement, leading to a map with an overall resolution of 3.34 Å, as determined by the Fourier shell correlation (FSC) = 0.143 criterion. The local resolution map was analyzed using Local Resolution Estimation of cryoSPARC and displayed by UCSF ChimeraX^24^.

The model of FcRL4 was predicted by Alphafold^25^. Models of Fcα–J complex was taken from the structure of the SIgA complex (PDB ID: 6LX3). Models and the cryo-EM map were fitted by UCSF chimeraX and manually adjusted using Coot^26^. The density map for the FcRL4-D4 domain was of insufficient quality for model building. Refinement was performed using the real-space refinement in Phenix^27^. Figures were prepared with UCSF ChimeraX.

### StrepTactin pull-down assay

To verify the interaction between Fcα–J and FcRL4, wild type and mutant FcRL4 proteins were purified and incubated with purified Fcα–J on ice for 1 h. The mixture was then incubated with the StrepTactin beads (Smart Lifesciences) and rotated at 4 °C, in the binding buffer containing 25 mM Tris-HCl, pH 7.4, 150 mM NaCl. Fcα features an N-terminal TwinStrep tag. After 1 h of incubation, the beads were spun down and washed with the binding buffer three times. The bound proteins were eluted with the binding buffer supplemented with 10 mM desthiobiotin (IBA Lifesciences). The samples were analyzed by SDS-PAGE and detected by immunoblotting using antibodies to the strep tag (1:3,000; A01736-100, GenScript) and His tag (1:1,500; HT501, TransGen Biotech).

## Data and materials availability

Cryo-EM density maps of Fcα–J–FcRL4 have been deposited in the Electron Microscopy Data Bank with accession codes EMD-37943. Structural coordinates have been deposited in the Protein Data Bank with the accession codes 8WZ1.

## Acknowledgments

We are grateful to the Core Facilities at the School of Life Sciences, Peking University for their assistance with negative-staining EM; the Cryo-EM Platform of Peking University for their support with data collection; and the High-performance Computing Platform of Peking University for their aid with computation. We also acknowledge the National Center for Protein Sciences at Peking University and Changping Laboratory for their help with Biacore facilities. This work received support from the National Natural Science Foundation of China (32325018) and the Qidong-SLS Innovation Fund to J.X., as well as from Changping Laboratory.

## Author Contributions

C.S. and Y.W. contributed equally to this work. C.S. performed the IgA binding assays and SPR analyses. C.S. and Y.W. carried out protein expression, purification, cryo-EM sample screening and data collection. Y.W. performed the cryo-EM data processing, model building and structure refinement with assistance from X.Y. C.S. and Y.W. conducted the pull-down experiments. J.X. conceived and supervised the project and wrote the manuscript, with inputs from all authors.

---

**Extended Data Fig. 1.**
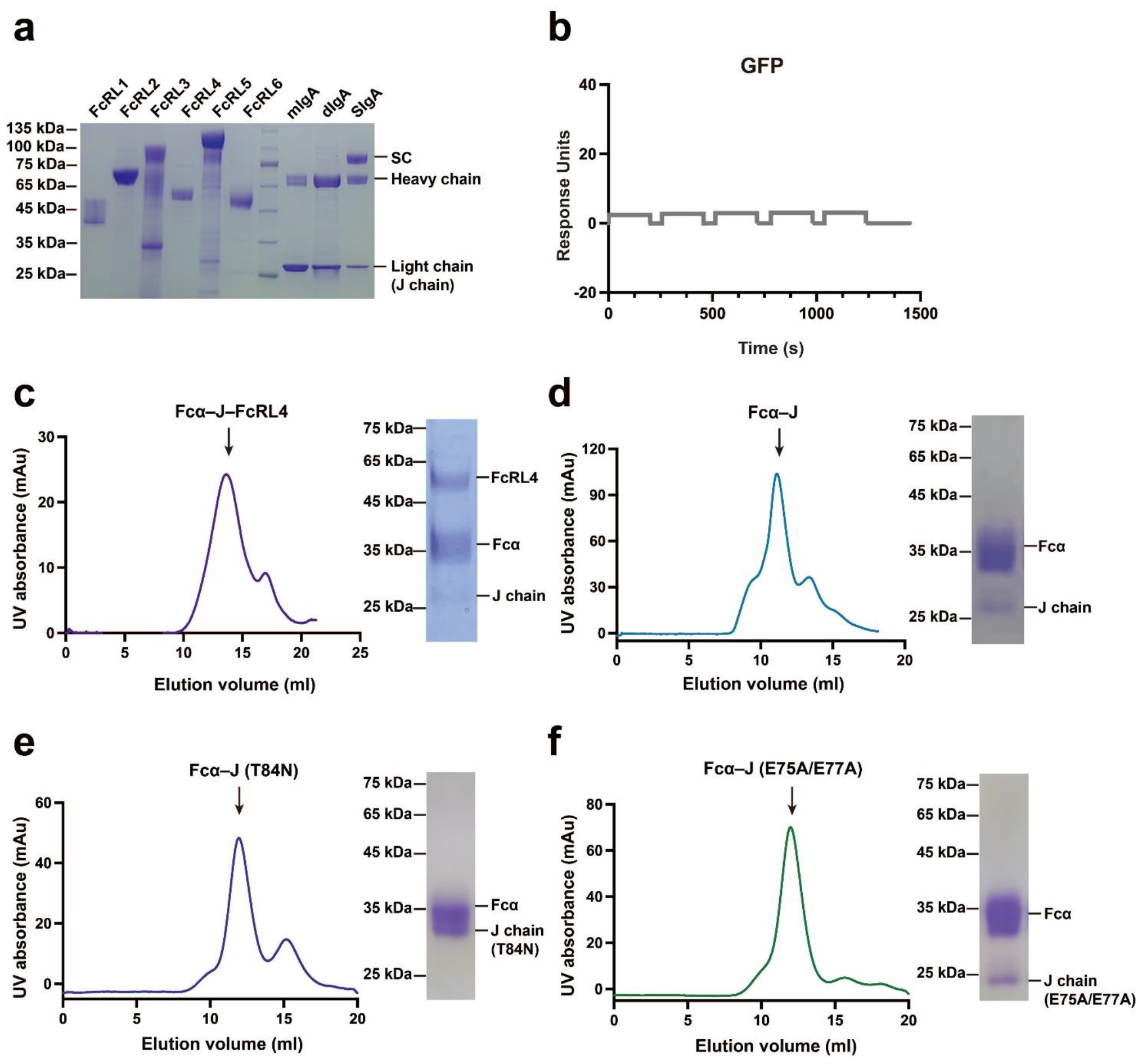
Purified proteins used in this study. **a**. SDS–PAGE analyses of the extracellular domains of human FcRL1–6 and various forms of IgA. The mIgA-604 sample was purified by anti-light chain affinity chromatography using Capto-L beads, and therefore contains extra light chains. **b**. Single cycle kinetics SPR analysis of the interaction between Fcα–J and immobilized GFP (negative control). **c**. Size-exclusion chromatography of the Fcα–J–FcRL4 complex on a Superose 6 Increase column. **d–f**. Size-exclusion chromatography profiles and SDS–PAGE analyses of the Fcα–J complexes.

**Extended Data Fig. 2.**
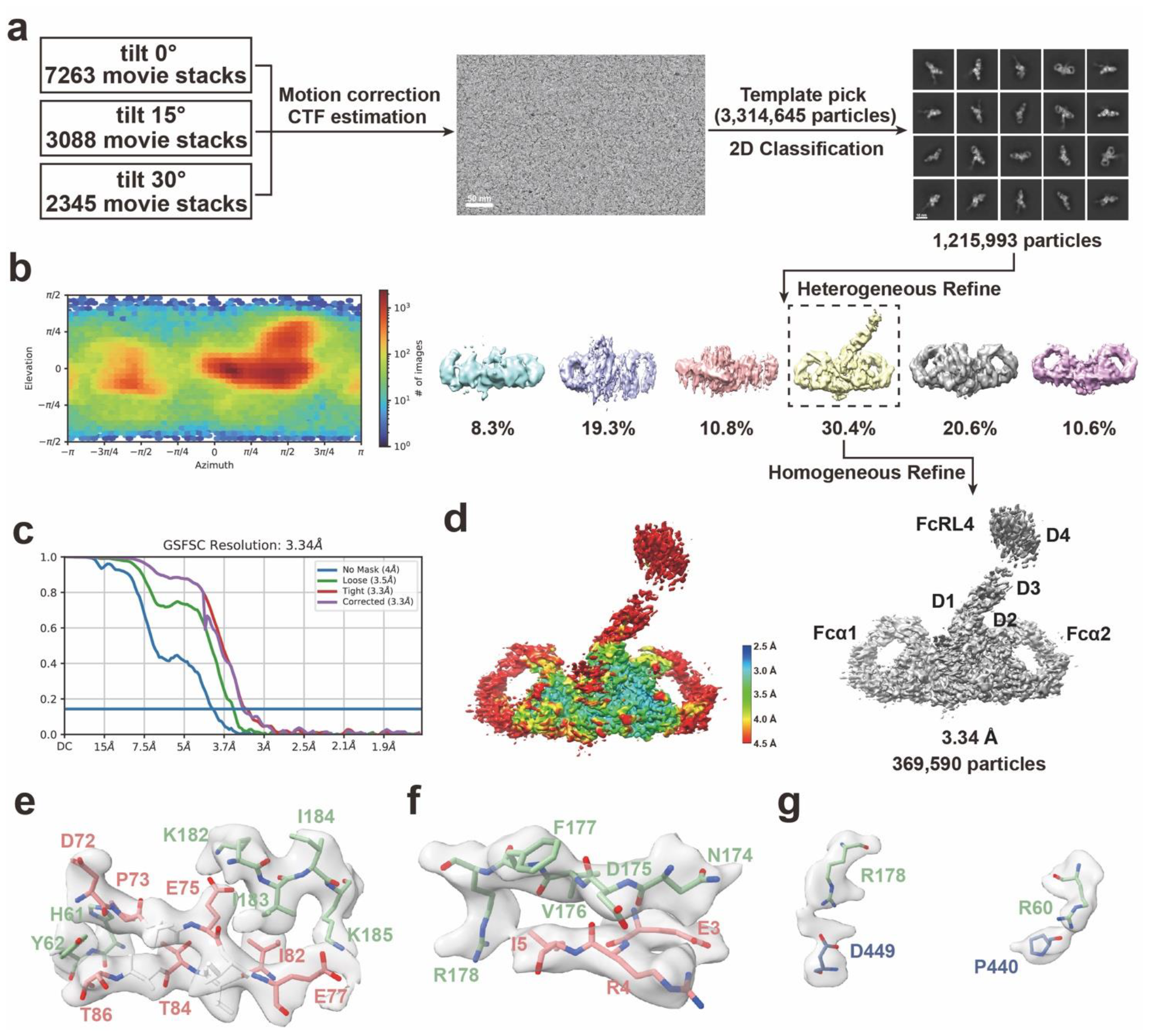
Cryo-EM structure determination of the Fcα–J–FcRL4 complex. **a**. Flowchart of cryo-EM data processing. **b**. Angular particle distribution heat map. **c**. Gold-standard Fourier shell correlation (GSFSC) curves of the Fcα–J–FcRL4 complex. **d**. Resolution estimations for the final maps of the Fcα–J–FcRL4 complex. **e**. Cryo-EM densities reveal that FcRL4 interacts with the β5–β6 hairpin of the J chain. **f**. Cryo-EM densities reveal that FcRL4 interacts with the N-terminal region of the J chain. **g**. Cryo-EM densities reveal that FcRL4 interacts with Fcα2.

**Extended Data Fig. 3.**
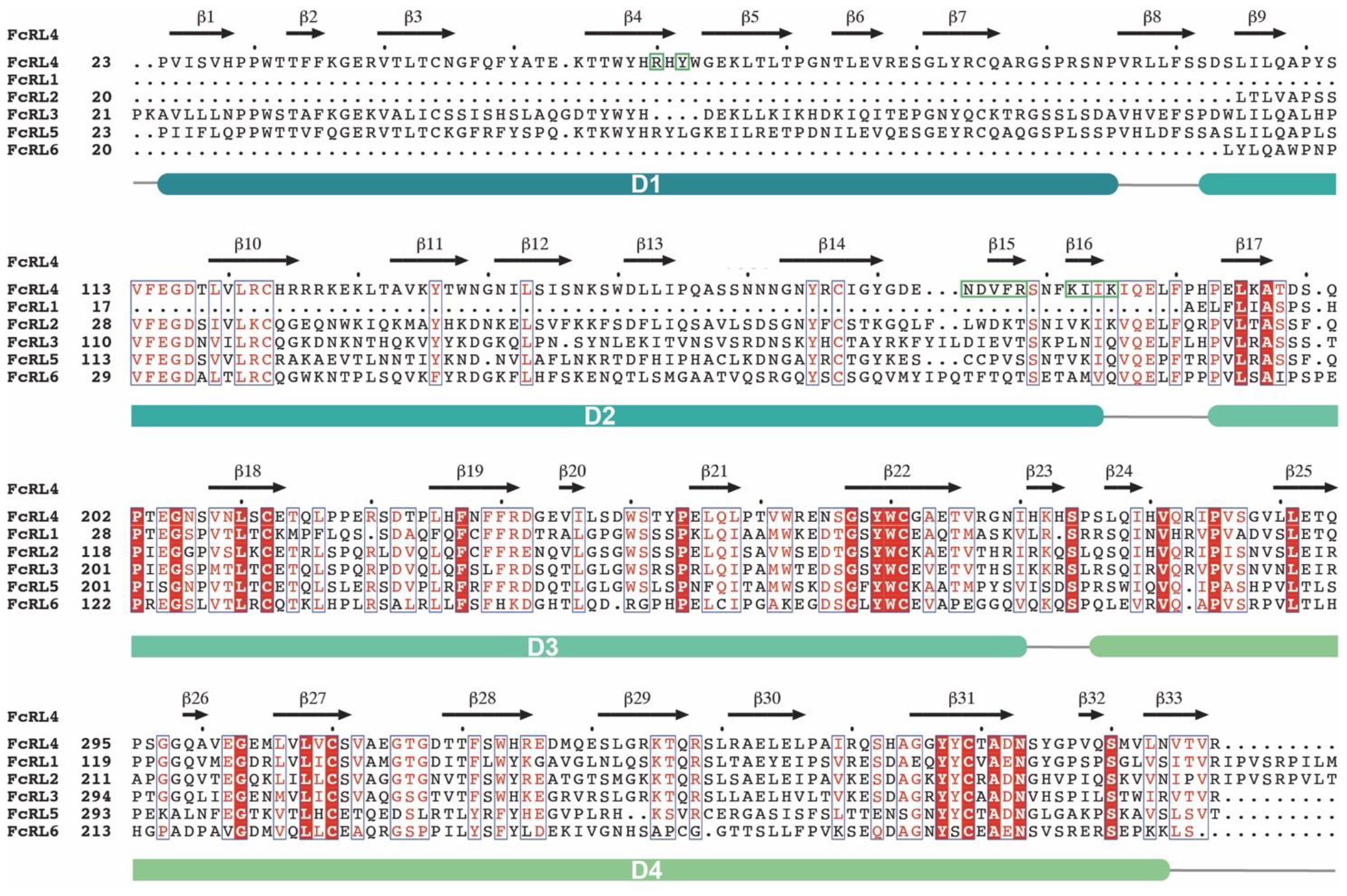
Sequence alignment of FcRL family members. Residues in human FcRL4 that are involved in binding to Fcα–J are highlighted in green boxes. Four Ig-like domains of FcRL4 are shown in teal, light teal, sea green and light sea green, respectively.

**Extended Data Fig. 4.**
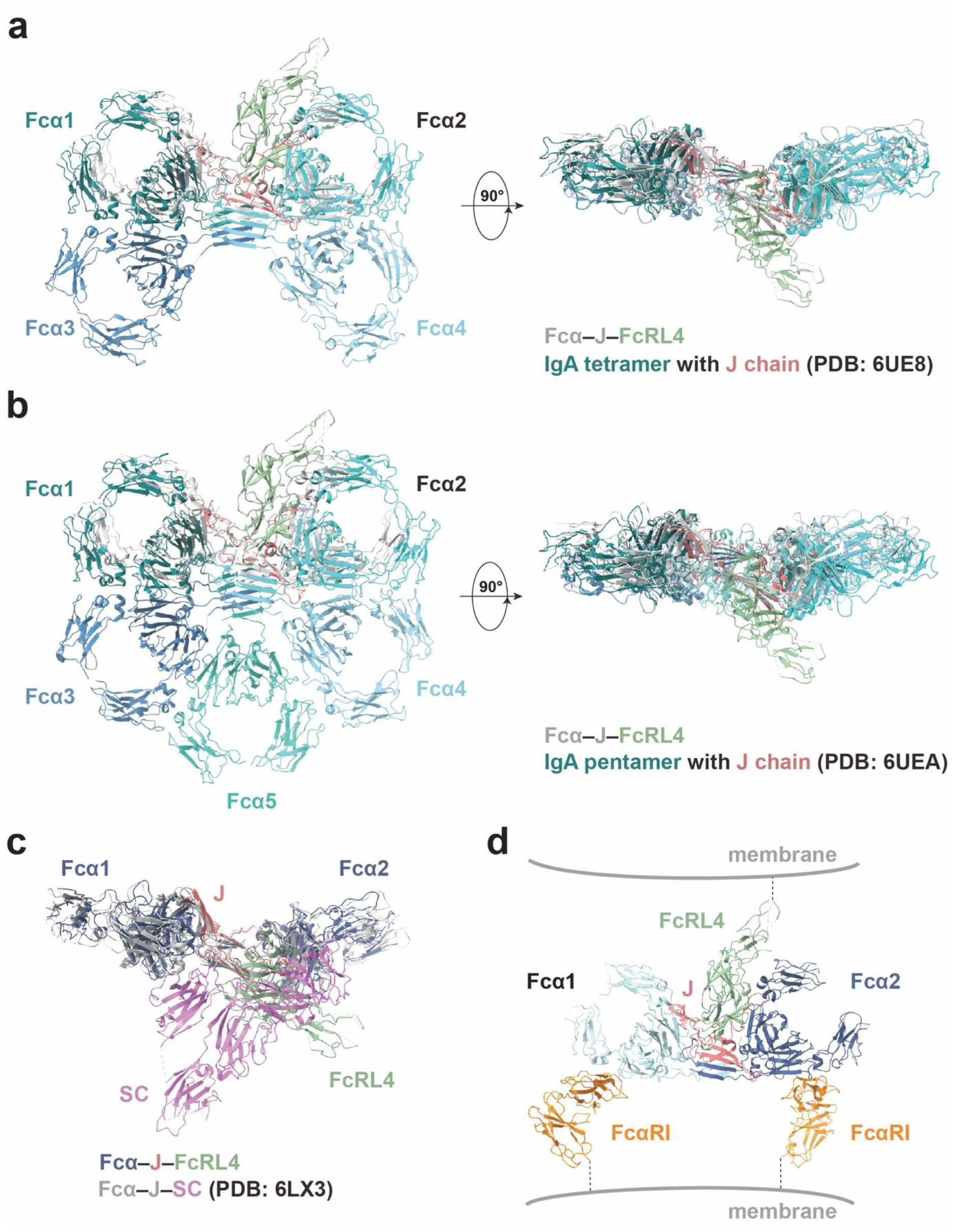
Structural relationship between FcRL4 and tetrameric IgA, pentameric IgA, secretory IgA, and FcαRI. **a**. FcRL4 should be able to bind to tetrameric IgA. **b**. FcRL4 should be able to bind to pentameric IgA as well. **c**. Structural analysis suggests that SC blocks the binding of FcRL4. **d**. A hypothetical model of the Fcα–J–FcRL4-FcαRI complex.

**Extended Data Table 1.** Cryo-EM data collection, refinement and validation statistics.

|  |  |
| --- | --- |
|  | IgA1-Fc/J/FcRL4 |
|  | PDB ID: 8WZ1 |
|  | EMDB: EMD-37943 |
| **Data collection and processing** |  |
| Magnification | 105,000 |
| Voltage (kV) | 300 |
| Electron exposure (e-/Å<sup>2</sup>) | 59.5 |
| Defocus range (µm) | −0.8 to −1.6 |
| Pixel size (Å) | 0.83 |
| Symmetry imposed | C1 |
| Initial particle images (no.) | 3,314,645 |
| Final particle images (no.) | 369,590 |
| Map resolution (Å) | 3.34 |
| FSC threshold: 0.143 |  |
| **Refinement** |  |
| Initial model used (PDB code) | 6LX3 |
| Model resolution (Å) | 3.34 |
| FSC threshold: 0.143 |  |
| Map sharpening *B* factor (Å<sup>2</sup>) | 101.1 |
| Model composition |  |
| Non-hydrogen atoms | 9,151 |
| Protein residues | 1,167 |
| Ligands | 8 |
| *B* factors (Å<sup>2</sup>) |  |
| Protein | 56.98 |
| Ligand | 89.44 |
| R.m.s. deviations |  |
| Bond lengths (Å) | 0.006 |
| Bond angles (°) | 1.063 |
| Validation |  |
| MolProbity score | 1.72 |
| Clashscore | 6.76 |
| Poor rotamers (%) | 1.08 |
| Ramachandran plot |  |
| Favored (%) | 95.31 |
| Allowed (%) | 4.69 |
| Disallowed (%) | 0.00 |

## Reference

1. Woof, J.M. & Kerr, M.A. The function of immunoglobulin A in immunity. J Pathol 208, 270–82 (2006).

2. Matsumoto, M.L. Molecular Mechanisms of Multimeric Assembly of IgM and IgA. Annu Rev Immunol 40, 221–247 (2022).

3. Monteiro, R.C. & Van De Winkel, J.G. IgA Fc receptors. Annu Rev Immunol 21, 177–204 (2003).

4. Wilson, T.J., Fuchs, A. & Colonna, M. Cutting edge: human FcRL4 and FcRL5 are receptors for IgA and IgG. J Immunol 188, 4741–5 (2012).

5. Liu, Y. et al. FCRL4 Is an Fc Receptor for Systemic IgA, but Not Mucosal Secretory IgA. J Immunol 205, 533–538 (2020).

6. Davis, R.S. Fc receptor-like molecules. Annu Rev Immunol 25, 525–60 (2007).

7. Hatzivassiliou, G. et al. IRTA1 and IRTA2, novel immunoglobulin superfamily receptors expressed in B cells and involved in chromosome 1q21 abnormalities in B cell malignancy. Immunity 14, 277–89 (2001).

8. Davis, R.S., Wang, Y.H., Kubagawa, H. & Cooper, M.D. Identification of a family of Fc receptor homologs with preferential B cell expression. Proc Natl Acad Sci U S A 98, 9772–7 (2001).

9. Delidakis, G., Kim, J.E., George, K. & Georgiou, G. Improving Antibody Therapeutics by Manipulating the Fc Domain: Immunological and Structural Considerations. Annu Rev Biomed Eng 24, 249–274 (2022).

10. Falini, B. et al. Expression of the IRTA1 receptor identifies intraepithelial and subepithelial marginal zone B cells of the mucosa-associated lymphoid tissue (MALT). Blood 102, 3684–92 (2003).

11. Ehrhardt, G.R. et al. Expression of the immunoregulatory molecule FcRH4 defines a distinctive tissue-based population of memory B cells. J Exp Med 202, 783–91 (2005).

12. Miller, I., Hatzivassiliou, G., Cattoretti, G., Mendelsohn, C. & Dalla-Favera, R. IRTAs: a new family of immunoglobulinlike receptors differentially expressed in B cells. Blood 99, 2662–9 (2002).

13. Ehrhardt, G.R. et al. The inhibitory potential of Fc receptor homolog 4 on memory B cells. Proc Natl Acad Sci U S A 100, 13489–94 (2003).

14. Sohn, H.W., Krueger, P.D., Davis, R.S. & Pierce, S.K. FcRL4 acts as an adaptive to innate molecular switch dampening BCR signaling and enhancing TLR signaling. Blood 118, 6332–41 (2011).

15. Yeo, L. et al. Expression of FcRL4 defines a pro-inflammatory, RANKL-producing B cell subset in rheumatoid arthritis. Ann Rheum Dis 74, 928–35 (2015).

16. Amara, K. et al. B cells expressing the IgA receptor FcRL4 participate in the autoimmune response in patients with rheumatoid arthritis. Journal of Autoimmunity 81, 34–43 (2017).

17. Du, S. et al. Structurally Resolved SARS-CoV-2 Antibody Shows High Efficacy in Severely Infected Hamsters and Provides a Potent Cocktail Pairing Strategy. Cell 183, 1013–1023 e13 (2020).

18. Agarwal, S. et al. Human Fc Receptor-like 3 Inhibits Regulatory T Cell Function and Binds Secretory IgA. Cell Rep 30, 1292–1299 e3 (2020).

19. Herr, A.B., White, C.L., Milburn, C., Wu, C. & Bjorkman, P.J. Bivalent binding of IgA1 to FcalphaRI suggests a mechanism for cytokine activation of IgA phagocytosis. J Mol Biol 327, 645–57 (2003).

20. Herr, A.B., Ballister, E.R. & Bjorkman, P.J. Insights into IgA-mediated immune responses from the crystal structures of human FcαRI and its complex with IgA1-Fc. Nature 423, 614–620 (2003).

21. Liu, Q. & Stadtmueller, B.M. SIgA structures bound to Streptococcus pyogenes M4 and human CD89 provide insights into host-pathogen interactions. Nat Commun 14, 6726 (2023).

22. Wang, Y.X. et al. Structural insights into secretory immunoglobulin A and its interaction with a pneumococcal adhesin. Cell Research 30, 602–609 (2020).

23. Punjani, A., Rubinstein, J.L., Fleet, D.J. & Brubaker, M.A. cryoSPARC: algorithms for rapid unsupervised cryo-EM structure determination. Nature methods 14, 290–296 (2017).

24. Pettersen, E.F. et al. UCSF ChimeraX: Structure visualization for researchers, educators, and developers. Protein Science 30, 70–82 (2021).

25. Jumper, J. et al. Highly accurate protein structure prediction with AlphaFold. Nature 596, 583–589 (2021).

26. Emsley, P., Lohkamp, B., Scott, W.G. & Cowtan, K. Features and development of Coot. Acta Crystallographica Section D: Biological Crystallography 66, 486–501 (2010).

27. Adams, P.D. et al. PHENIX: a comprehensive Python-based system for macromolecular structure solution. Acta Crystallographica Section D: Biological Crystallography 66, 213–221 (2010).

